# EGFR does not directly interact with cortical actin: A SRRF’n’TIRF Study

**DOI:** 10.1101/2024.07.20.604398

**Authors:** Shambhavi Pandey, Thorsten Wohland

## Abstract

The epidermal growth factor receptor (EGFR) governs pivotal signaling pathways in cell proliferation and survival, with mutations implicated in numerous cancers. The organization of EGFR on the plasma membrane (PM) is influenced by the lipids and the cortical actin (CA) cytoskeleton. Despite the presence of a putative actin-binding domain (ABD) spanning 13 residues, a direct interaction between EGFR and CA has not been definitively established. While disrupting the cytoskeleton can impact EGFR behavior, suggesting a connection, the influence of the static actin cytoskeleton has been found to be indirect. Here, we investigate the potential interaction between EGFR and CA, as well as the extent to which CA regulates EGFR’s distribution on the PM using SRRF’n’TIRF, a spatiotemporal super-resolution microscopy technique that provides sub-100 nm resolution and ms-scale dynamics from the same dataset. To label CA, we constructed PMT-mEGFP-F-tractin, which combines an inner leaflet targeting domain PMT, fluorescent probe mEGFP, and the actin-binding protein F-tractin. In addition to EGFR-mEGFP, we included two control constructs: a) an ABD deletion mutant, EGFR^ΔABD^-mEGFP serving as a negative control, and b) EGFR-mApple-F-tractin, where F-tractin is fused to the C-terminus of EGFR-mApple, serving as the positive control. We find that EGFR-mEGFP and EGFR^ΔABD^-mEGFP show similar membrane dynamics, implying that EGFR-mEGFP dynamics and organization are independent of CA. EGFR dynamics show CA dependence when F-tractin is anchored to the cytoplasmic tail. Together, our results demonstrate that EGFR does not directly interact with the CA in its resting and activated state.

**SIGNIFICANCE:** SRRF’n’TIRF is a spatiotemporal super-resolution microscopy technique that allows for the investigation of plasma membrane-cytoskeleton interactions. We investigate how cortical actin (CA) influences the dynamic behavior and structural organization of EGFR, employing specific probe targeting CA structure and dynamics. Our results suggest that EGFR, whether in its resting or activated state, does not directly bind to or interact with the CA. Any influence of CA on EGFR is indirect through membrane modulating activities of CA.

## INTRODUCTION

The Epidermal Growth Factor Receptor (EGFR) is a trans-membrane glycoprotein that belongs to the family of receptor tyrosine kinases (RTKs) and is a key player in regulating cell proliferation, differentiation, survival, function, and motility (1–3). Structurally, EGFR comprises an extracellular domain, a hydrophobic membrane-spanning region, and an intracellular tyrosine kinase domain, and the process of its activation by ligands (such as betacellulin, amphiregulin (AR), epidermal growth factor (EGF), heparin-binding EGF-like growth factor, and transforming growth factor (TGF-*α*)) has been extensively researched (4–6). Various studies have supported the model wherein resting state EGFR exists as monomers in equilibrium with a pool of inactive dimers (7, 8) (Figure 1a). When the ligands bind to EGFR, they stabilize receptor conformations, exposing the dimerization interface on the extracellular side. This event triggers the accumulation of active EGFR dimers, leading to oligomerization (9, 10). Upon activation, EGFR undergoes autophosphorylation on five tyrosine residues located within its C-terminal regulatory domain. These phosphorylated tyrosine residues then act as binding sites for downstream signaling molecules, initiating a cascade of downstream signaling pathways (11, 12). Overexpression and/or elevation in EGFR’s activity contributes to the progression of human cancers. EGFR has been found to be overexpressed in various solid tumors, such as breast, head and neck (H&N), colon, ovarian, renal, pancreatic, and non-small-cell lung cancer (3, 13–17). These cancers are characterized by highly metastatic, aggressive, and drug-resistant tumors (18). Thus, EGFR is a promising target for developing anticancer therapeutics (19–21).

**Figure 1:**
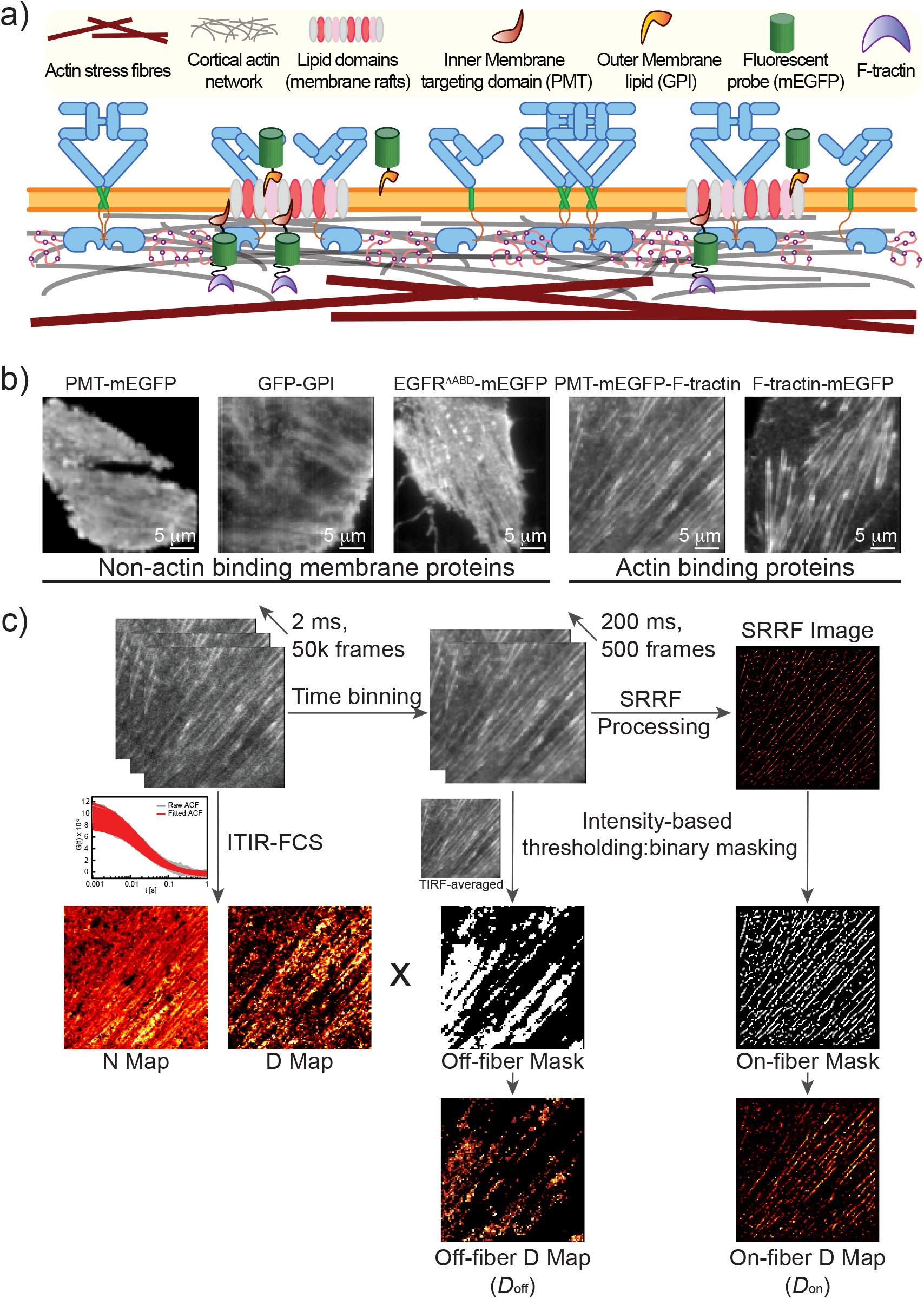
Membrane Schematics and SRRF’n’TIRF FCS Workflow. a) Membrane schematics illustrate the distribution of EGFR on the membrane, ranging from monomers to oligomers, as well as the presence of lipid domains and various outer (GPI) and inner (PMT) leaflet lipids. The dynamic CA network (grey closely apposes the membrane, while long stress fibers (brown) are situated further away. b) TIRF-averaged images reveal actin fiber structures, observed with both actin-binding membrane proteins and non-actin binding lipids/proteins. c) The SRRF’n’TIRF FCS workflow enables simultaneous investigation of dynamics (ITIR-FCS) and structure (SRRF) using the same dataset. SRRF reconstruction of the TIRF stack allows for the masking of on-fiber regions, while the TIRF-averaged image was used to create the off-fiber mask, which was subsequently used to generate on-/off-fiber D maps.

EGFR is a highly regulated molecule, with its regulation influenced by its inherent structural features and the composition of the plasma membrane (PM) environment. The PM consists of a phospholipid bilayer containing various lipids, primarily classified as phospholipids, glycolipids, and sterols (22). As an embedded transmembrane receptor, EGFR interacts with lipids in both the outer and inner leaflets of the PM. These interactions are crucial for EGFR activation and have been extensively investigated (23–25). Recent studies have identified several lipids involved in these interactions, including phosphatidylcholine (PC), phosphatidylserine (PS), phosphatidylinositol 4,5-bisphosphate (PI(4,5)P2), cholesterol, gangliosides, and palmitate (26–33). Membrane lipids are integral to maintaining the biophysical properties of the PM (22) and are known to form transient lipid-ordered domains, also known as membrane (lipid) rafts (Figure 1a). These rafts are dynamic, laterally heterogeneous, and enriched with cholesterol or sphingolipids (34, 35). EGFR can form higher-order oligomers (36, 37), and these clusters are sensitive to both activating ligand dosage and the lipid-ordered domains on the PM (38, 39). Notably, about 50-70% of the resting EGFR exist as preformed dimers or oligomers (40–45). The lipid-ordered domains play a crucial role in retaining portions of the EGFR population and preventing unintended clustering and activation of the receptor in its resting state. Disrupting these domains by depleting cholesterol or removing sphingolipids enhances EGFR clustering, facilitating its activation in both ligand-dependent and ligand-independent manners (29, 39).

Apart from membrane lipids, it is well established that the cortical actin (CA) - the isotropic actin network bound and located proximally to the PM - significantly influences membrane organization. Acting as a dynamic scaffold, CA provides structural support and mechanical force for PM remodeling and receptor organization and dynamics (46, 47). Local changes in CA density can affect the diffusion mode and rate of membrane lipids and proteins, as well as membrane tension (48–50). However, due to its lower density compared to stress fibers, which are primarily found deeper within the cell and form linear arrays of actin filaments, the presence of the CA network in fluorescence images of the actin cytoskeleton can be easily overlooked (51). The complex interplay between the PM and CA architecture complicates the study of these links, highlighting the need for techniques that allow simultaneous quantification of membrane components’ dynamics and CA structure.

In this context, the influence of the CA network on EGFR dynamics and organization remains poorly understood. A region within the intracellular kinase domain of EGFR, spanning amino acid residues 984-996, forms an actin-binding domain (ABD) (52), with documented associations with the cytoskeleton (53–56). Disruptions to the actin cytoskeleton can alter EGFR behavior, suggesting a connection (39). Recent work from our group has shown that the actin cytoskeleton indirectly influences EGFR dynamics and organization by altering membrane properties (57). However, the actin cytoskeleton signal could not be conclusively identified as originating from CA or stress fibers, leaving the precise CA-EGFR interaction unresolved. Despite the presence of a putative ABD, a direct interaction between EGFR and CA has not yet been established, and links between the resting/activated-state EGFR and the dynamic CA are still an open question. Further investigations are required to elucidate the precise nature of this interaction and whether it entails a direct or indirect connection.

Here, we use SRRF’n’TIRF (58), combining imaging total internal reflection fluorescence correlation spectroscopy (ITIR-FCS) and super-resolution radial fluctuations (SRRF) to study the structure and dynamics in live CHO-K1 cells. TIRF data allows for correlating spatial and temporal fluctuations, with a single TIRF image stack used to extract both structural and dynamics information. Dynamics are quantified using ITIR-FCS. This method employs TIR illumination to excite fluorescent molecules, with a fast, sensitive camera capturing the fluorescence signal (58–60). Unlike conventional FCS, ITIR-FCS simultaneously analyzes fluorescence fluctuations at thousands of contiguous spots, generating spatially resolved parametric maps. To enhance structural details, we apply SRRF (61), a computational technique using radial intensity gradients to improve resolution beyond the optical diffraction limit. SRRF’n’TIRF achieves a spatial resolution of less than 100 nm and captures dynamics on the millisecond timescale, enabling the simultaneous study of CA structure and membrane protein dynamics from a single measurement.

We specifically labeled the CA, using PMT-mEGFP/mApple-F-tractin (Figure 1a), a probe that comprises an inner leaflet PM targeting peptide (PMT), a fluorescent probe (mEGFP/mApple), and an F-actin-binding protein (F-tractin), inspired by (62). We generated control constructs of EGFR, one lacking its putative actin-binding domain, EGFR^ΔABD^-mEGFP as the negative control, and the other where F-tractin is anchored to its cytoplasmic tail, EGFR-mApple-F-tractin as the positive control. Our results show that the diffusion of EGFR-mEGFP and EGFR^ΔABD^-mEGFP are not influenced by the CA, unlike PMT-mEGFP-F-tractin and EGFR-mApple-F-tractin, which showed reduced diffusion at locations of higher CA density, implying that EGFR does not directly interact with the CA in its resting or activated state.

## MATERIALS AND METHODS

### Cell Culture

CHO-K1 cells (CCL-61™, ATCC; Manassas, Virginia, USA) were cultivated in Dulbecco’s Modified Eagle Medium (DMEM/High glucose with L-glutamine, excluding sodium pyruvate – SH30022.FS, HyClone, GE Healthcare Life Sciences, Utah, USA). The medium was supplemented with 1% penicillin-streptomycin (#15070063, Gibco, Thermo Fisher Scientific, Massachusetts, USA) and 10% fetal bovine serum (FBS; #10270106, Gibco). The cells were cultured in 25 cm^2^ polystyrene cell culture flasks (#430639, Corning, New York, USA), and regular passages were conducted upon reaching confluency. The cell cultures were maintained within a controlled environment of a heated CO2 incubator at 37°C and 5% (v/v) CO2 concentration (Forma Steri-Cycle CO2 incubator, Thermo Fisher Scientific).

### Plasmid Construction

The construction of PMT-mEGFP (Addgene #203777)/-mApple (Addgene #203771) and EGFR-mEGFP (Addgene #2037714/- mApple (Addgene #203773) has been detailed elsewhere (57). The GFP-GPI plasmid was a gift from Dr. John Dangerfield (Anovasia Pte Ltd, Singapore). Recombinant PMT-mEGFP-F-tractin and PMT-mApple-F-tractin plasmids were created using PMT-mEGFP/PMT-mApple as a DNA template and adding F-tractin sequence by PCR with forward primer flanked by restriction enzymes NheI and reverse primer flanked by HindIII. Subsequently, PMT sequences were removed by AgeI/SpeI restriction enzyme digest to generate F-tractin-mEGFP/-mApple plasmids. The EGFR-mApple-F-tractin construct involved inserting the mApple-F-tractin sequence in-frame at the 3’ end of the EGFR cytoplasmic domain (without the stop codon). The amplicon containing F-tractin was PCR amplified from the PMT-mApple-F-tractin plasmid, and the overlapping end sequence of EGFR-mApple was PCR amplified from the EGFR-mApple plasmid using appropriate primer pairs. The recombinant EGFR^ΔABD^ construct featured a deletion of 13 amino acids (DDVVDADEYLIPQ) in the EGFR cytoplasmic domain of EGFR-mEGFP. The amplicon introducing this 39 bp deletion was PCR amplified from the EGFR-mEGFP plasmid, and the overlapping EGFR-mEGFP end sequence was generated using specific primer pairs.

All PCR amplification was conducted using Q5 High-Fidelity DNA Polymerase 2x master mix (M0492L, NEB). Gibson assembly of purified DNA fragments was carried out using NEBuilder HiFi DNA Assembly Master Mix (E2621L, NEB). NEB 5-alpha Competent E. coli cells (C2987H, NEB) were used to isolate respective recombinant clones.

### Cell Transfection

Cell cultures at approximately 90% confluence were utilized for transfection. The spent media was aspirated, and the culture flask underwent two washes with 5 ml of 1X PBS (phosphate-buffered saline lacking Ca2+ and Mg2+). Next, 2 ml of Trypsin-EDTA (0.5%; #15400054, Gibco) was introduced, and the flask was incubated at 37°C for 2-3 minutes to facilitate cell detachment. To stop trypsin activity, 5 ml of culture media was added, and the cell suspension was centrifuged at 200g for 3 minutes. After discarding the supernatant, the cell pellet was resuspended in 5 ml of PBS. The cell count was performed using an automated cell counter (#TC20, Bio-Rad, California, USA). The requisite number of cells (2− 5 × 10^5^) was pelleted by centrifugation (#5810, Eppendorf, Hamburg, Germany) at 200g for 3 minutes and the supernatant was discarded. The cell pellet was then resuspended in R buffer (Neon Transfection Kit, Thermo Fisher). Appropriate amounts of plasmids (100 ng for the PMT-F-tractin/PMT/GPI/F-tractin plasmids; 1 µg for the EGFR plasmids) were mixed with the cells. The transfection of cells was carried out using the lipid-based transfection reagent (Lipofectamine™ 3000, Invitrogen™), adhering to the manufacturer’s protocol. The transfected cells were then seeded onto culture dishes (#P35G-1.5-20-C, MatTek, Massachusetts, USA) containing DMEM (supplemented with 10% FBS; no antibiotics) and incubated at 37°C in a 5% CO2 environment for 36–48 hours before performing measurements.

### Sample Preparation for Fluorescence Measurements

The transfected cells underwent a rinse with Hank’s Balanced Salt Solution (1X HBSS with Ca2+ and Mg2+; #14025134, Gibco), followed by the addition of DMEM lacking phenol red (#21063029, Gibco). For the work presented in this article, DMEM without phenol red is called “Imaging DMEM”.

### Ligand Stimulation and Drug Treatments

Working concentrations of the drugs were prepared using imaging DMEM. For ligand stimulation, cells were treated with 100 ng/mL hEGF (#E9644, Sigma-Aldrich, Singapore) for 15 minutes. To achieve actin cytoskeleton polymerization and disruption, cells were treated with 1 µM Jasplakinolide (Jas, #J7473, Sigma-Aldrich) and 1 µM Latrunculin-A (Lat-A, #L5163, Sigma-Aldrich) for 30 minutes each, respectively. For stress fiber inhibition, cells were plated on poly-L-lysine (#P4832, Sigma-Aldrich) coated dishes and cultured with 1 µM Rho-associated kinase inhibitor Y-27632 (#Y0503, Sigma-Aldrich) mixed media for 24 hours before transfection and 48 hours before measurements.

### Confocal Imaging

Z-stack confocal imaging was performed using a commercial Zeiss LSM 900 microscope equipped with an Airyscan II detector (Carl Zeiss, Oberkochen, Germany). The 488 nm and 561 nm lasers were used to illuminate mEGFP-tagged and mApple-tagged constructs, respectively. Images were captured at a resolution of 1024 by 1024 pixels using a 63x (NA 1.4) objective with a z-step size of 0.28 µm. Images were saved in the “czi” format, and analysis was performed using Imaris 9.6.0 (Bitplane, South Windsor, CT, USA) and Zen Blue 3.1 (Carl Zeiss) softwares.

### Instrumentation

The TIRF microscopy setup includes an inverted epi-fluorescence microscope (IX83; Olympus, Singapore) with a PlanApo objective (100×, NA 1.49; Olympus) and a back-illuminated EMCCD camera (Andor iXON 860, 128×128 pixels; Andor Technology, US). A 488 nm (LAS/488/100, Olympus) and a 561 nm (LAS/561/100, Olympus) laser were coupled via the TIRF illumination combiner (cell TIRF/IX3-MITICO, Olympus) to the microscope. Cell measurements were conducted at 37°C and 5% CO2 using an on-stage incubator (Chamlide TC, Live Cell Instrument, South Korea).

A ZT 405/488/561/640rpc (Chroma Technology Corp, Vermont, USA) dichroic mirror was used to reflect the lasers onto the back focal plane of the objective, with the incidence angle adjusted using Olympus cellSens Imaging software to achieve TIRF mode. The signal, after passing through the objective and dichroic, was filtered using a laser quad band ZET405/488/561/647 m (Chroma Technology Corp) emission filter for TIRF applications and directed to the detector. For dual-channel measurements, a dual-emission image splitter (OptoSplit II; Cairn Research, Faversham, UK) was used, and the fluorescence light passed through a dichroic (ZT 405/488/561/640rpc, Chroma Technology Corp) and emission filter (ZET405/488/561/640 m, Chroma Technology Corp), before being split by the image splitter onto two halves of the camera chip. The image splitter was equipped with an emission dichroic (FF560-FDi01; Semrock, New York, USA) and band-pass filters (510AF23 and 585ALP, respectively; Omega Optical, Vermont, USA). A bright-field image of a stage micrometer was utilized for aligning the image splitter, carried out in µManager (version 1.4.14, µManager), and the alignment was completed by following the manual instructions.

### Data Acquisition

We used a laser power ranging from 100 µW to 1 mW (measured at the back focal plane of the objective) depending on the cell’s initial intensity count and expression level. The chosen penetration depth was 80 nm unless stated otherwise. Image acquisition was performed using Andor Solis software (v.4.31.30037.0-64-bit). The EMCCD camera was maintained at -80°C. The image acquisition used the kinetic mode, with the ‘baseline clamp’ consistently applied to minimize baseline fluctuations. The camera operated at a 10 MHz pixel readout speed, with the maximum analog-to-digital gain set to 4.7 and a vertical shift speed of 0.45 µs. An EM gain of 300 was used. We captured a stack of 50,000 frames, each with dimensions of 128 × 128 pixels, at a rate of 500 frames per second (2.06 ms per frame). These image stacks were saved in 16-bit TIFF format.

### ITIR-FCS Analysis

The image stacks from the cell measurements were loaded into the ImFCS 1.612 (63) plugin in Fiji. The source code for this plugin is available online (64). The parameters used were: frame time = 0.002 s, pixel size = 24 µm, NA = 1.49, *λ*1 = 507 nm (green channel)and *λ*1 = 565 nm (red channel). Correlator (p, q) = (16, 9) for on-fiber pixels and correlator (p,q) = (16,7) for off-fiber pixels for F-tractin constructs. For all other constructs, correlator (p, q) = (16, 9) was used. EMCCD data analysis was conducted at 1 × 1 binning with a ×100 magnification. Bleach correction (65) was performed using a polynomial of order 6, and the ACFs were fitted with a one-component diffusion model (58).Temporal binning of the entire stack was performed to obtain a TIRF-averaged image. Off-fiber pixels were selected using intensity thresholding, and an off-fiber binary mask was created in Fiji.

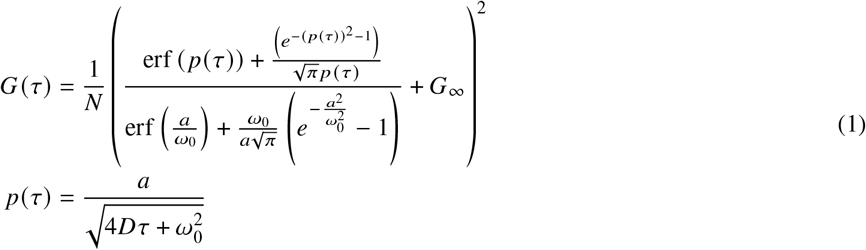

The provided equation *G* (*τ*) represents the theoretical model of the autocorrelation function (ACF), describing how it changes with the correlation time *τ* while recording the two-dimensional diffusion process. Parameters in the equation include ‘a’, which is the length of one pixel on the CCD chip, and *ω*_0_, representing the Gaussian approximation of the microscope’s point spread function. Additionally, *G*_∞_ signifies the ACF’s convergence value at long correlation times. The pixel side length is 24 µm, translating to a pixel side length, *a*, of 240 nm in the sample plane. *ω*_0_ is obtained by following the protocol described in (66). *N, D*, and *G*_∞_ were used as fit parameters, and the goodness-of-fit is assessed by *χ*^2^ value of the fit.

### SRRF Analysis

The acquired frame image stack was temporally binned in Fiji to a 206 ms, 500-frame stack (average binning of 100 frames each). The NanoJ-SRRF v1.14 plugin (61) was used with default parameter settings (ring radius 0.5, radiality magnification 5x, axes in the ring 6) and Temporal Radiality Auto-Correlations (TRAC) of order 2 for temporal analysis. The “display mode” was set to “radiality.” SRRF images were visually inspected for noise, granularity, patterns, or other features indicating artifacts. Intensity thresholding was applied to remove artifacts and select the on-fiber regions, and an on-fiber binary mask was generated in Fiji.

### Statistical Analysis

Results are expressed as mean ± standard deviation (SD). Data were statistically analyzed using two-way analysis of variance (ANOVA) in Python, followed by post-hoc tests to determine significance. Statistical significance among different groups is indicated as *** p < 0.001, * p < 0.05, and n.s. for non-significant differences.

## RESULTS AND DISCUSSION

### SRRF’n’TIRF FCS experimental protocol: On/Off dynamics

TIRF microscopy effectively minimizes background from deeper cell layers, providing high contrast and signal-to-noise ratio (SNR) for visualizing dynamic processes near the basal membrane of live cells. This selective optical sectioning is achieved by the evanescent wave generated at the glass-cell interface (67). In our imaging of membrane proteins, including actin-binding proteins (PMT-mEGFP-F-tractin, F-tractin-mEGFP) and non-actin-binding proteins (outer leaflet lipid glycosylphosphatidylinositol (GFP-GPI), inner leaflet lipid PMT-mEGFP and EGFR^ΔABD^-mEGFP), we observed actin fibers in the TIRF-averaged images (Figure 1b). Observing actin fibers with non-actin-binding proteins is likely attributed to membrane-modulating effects induced by actin at the cortex. Actin filaments exert force on the membrane, causing protrusions via various mechanisms (68). Additionally, several proteins such as the Arp2/3 complex, FBP17, and BAR domain proteins couple actin dynamics with membrane curvature (69–71). The intricate relationship between actin remodeling and membrane morphology, as discussed in recent review articles (72, 73), possibly explains the visualization of actin fiber-like structures in TIRF-averaged images of non-actin-binding proteins. However, the contrast of fibers for non-acting-binding proteins is very low compared to actin-binding proteins, and the two can be readily distinguished. We observed a gradual loss in contrast of fibers when imaging non-actin-binding protein, GFP-GPI (Figure S1a) as the penetration depth increased from 70 nm to 140 nm, while the fibers were retained when imaging F-tractin-mEGFP (Figure S1b) or PMT-mEGFP-F-tractin (Figure S1c). The fibers retained by PMT-mEGFP-F-tractin (Figure S1c) at a higher penetration depth of 140 nm indicate that the membrane-proximal fibers labeled by PMT-mEGFP-F-tractin dominate the SNR even at a penetration depth of 140 nm which is not the case with GFP-GPI. Simultaneous dual-wavelength measurements on cells expressing GFP-GPI and PMT-mApple-F-tractin confirmed that the fibers seen in the GFP-GPI channel are labeled by PMT-mApple-F-tractin (Figure S2a). Spatial intensity correlation of the fiber regions from the two images shows a similar correlation pattern along the fiber for both GFP-GPI and PMT-mApple-F-tractin (Figure S2b). This suggests that the actin-fiber-like structures seen for non-actin-binding proteins are indeed a membrane-related effect, and the observed fibers are actual actin fibers. In this study, we utilized the terms “on-fiber” for regions with observed fiber structures and “off-fiber” for regions devoid of fibers.

The schematic in Figure 1c illustrates the SRRF ‘n’ TIRF workflow employed in this study. The acquired TIRF image stack was analyzed using the standard ITIR-FCS protocol (74) as mentioned in the material and methods by acquiring 50,000 frames at 2.06 ms exposure per frame. The data was then used for FCS to calculate the autocorrelation function (ACF) for each pixel. FCS provides two key read-outs: the width of the ACF to determine the diffusion coefficient (D) and the amplitude of the ACF, which reflects the average number of particles (N) in the observation volume, which is related to concentration. The relationship between amplitude and concentration is dependent on the SNR and might need corrections if absolute values are to be determined (75, 76). These parameters are visualized as D and N maps, respectively. This stack is then time-binned either entirely to produce a time-binned TIRF-averaged image or partially by 100 frames, resulting in 500 frames of 206 ms exposure per frame for further processing to obtain a super-resolved image using SRRF. This allowed us to derive spatial information and temporal correlations from the same dataset, providing insights into both structure and dynamics. TIRF-averaged images are diffraction-limited, meaning pixels around fibers contain a mix of signals from both on-fiber and off-fiber regions, blurring the precise location of on-fiber pixels. However, the SRRF reconstruction, serving as the super-resolved image of the fibers, precisely identifies on-fiber pixels, which helps circumvent the issue of locating on-fiber pixels in the TIRF-averaged image. We used the SRRF image to create a binary mask for the on-fiber regions, while the diffraction-limited TIRF-averaged image was used to identify the off-fiber regions. Note that this approach identifies pixels containing clear fiber signals and those completely devoid of fibers while also excluding pixels adjacent to fibers that exhibit mixed on- and off-fiber signals. These on- and off-fiber binary masks were then employed to isolate on- and off-fiber D Maps from the original D Map. Subsequently, these maps were used to derive respective D-distributions, enabling us to observe the dynamics of the proteins in the on-/off-fiber regions and assess CA influence on protein dynamics and organization.

### Validation of PMT-F-tractin as the CA-binding structural and dynamic Control

Z-stack confocal imaging of CHO-K1 cells co-transfected with F-tractin-mApple and PMT-mEGFP-F-tractin shows distinctive localization patterns (Figure 2a). In the F-tractin-mApple channel, specific fibers are visible, but they are absent in the PMT-mEGFP-F-tractin channel, as observed in the Maximum Intensity Projection (MIP) images. This absence suggests that these fibers represent stress fibers located deeper within the cell, away from the membrane. The merged overlay shows distinct labeling patterns, with F-tractin-mApple distributed throughout the cell, while PMT-mEGFP-F-tractin primarily marks shorter dynamic CA filaments near the membrane. Localization of PMT-mEGFP-F-tractin and F-tractin-mApple at various depths within the cell shows PMT-mEGFP-F-tractin’s preference for labeling CA. However, it should be noted that some stress fiber-like structures can also be found close to the membrane and are thus labeled by both PMT-mEGFP-F-tractin and F-tractin-mApple. Subsequently, SRRF’n’TIRF analysis on cells expressing PMT-mEGFP-F-tractin demonstrated a distinct D-distribution pattern in the on-/off-fiber regions. The normalized frequency distribution of the on- and off-fiber D pooled from cells reveals a shift in D for off-fiber regions (0.35 ± 0.06 µm^2^ s^−1^) compared to the on-fiber areas (0.23 ± 0.06 µm^2^ s^−1^) (Figure 2b). The lower D of PMT-mEGFP-F-tractin in regions with fibers results from transient binding events that slow down PMT-mEGFP-F-tractin’s movements, similar to what has been seen for Lifeact previously (57, 77). Confocal and TIRF microscopy thus demonstrate the utility of PMT-F-tractin as a CA-binding probe.

**Figure 2:**
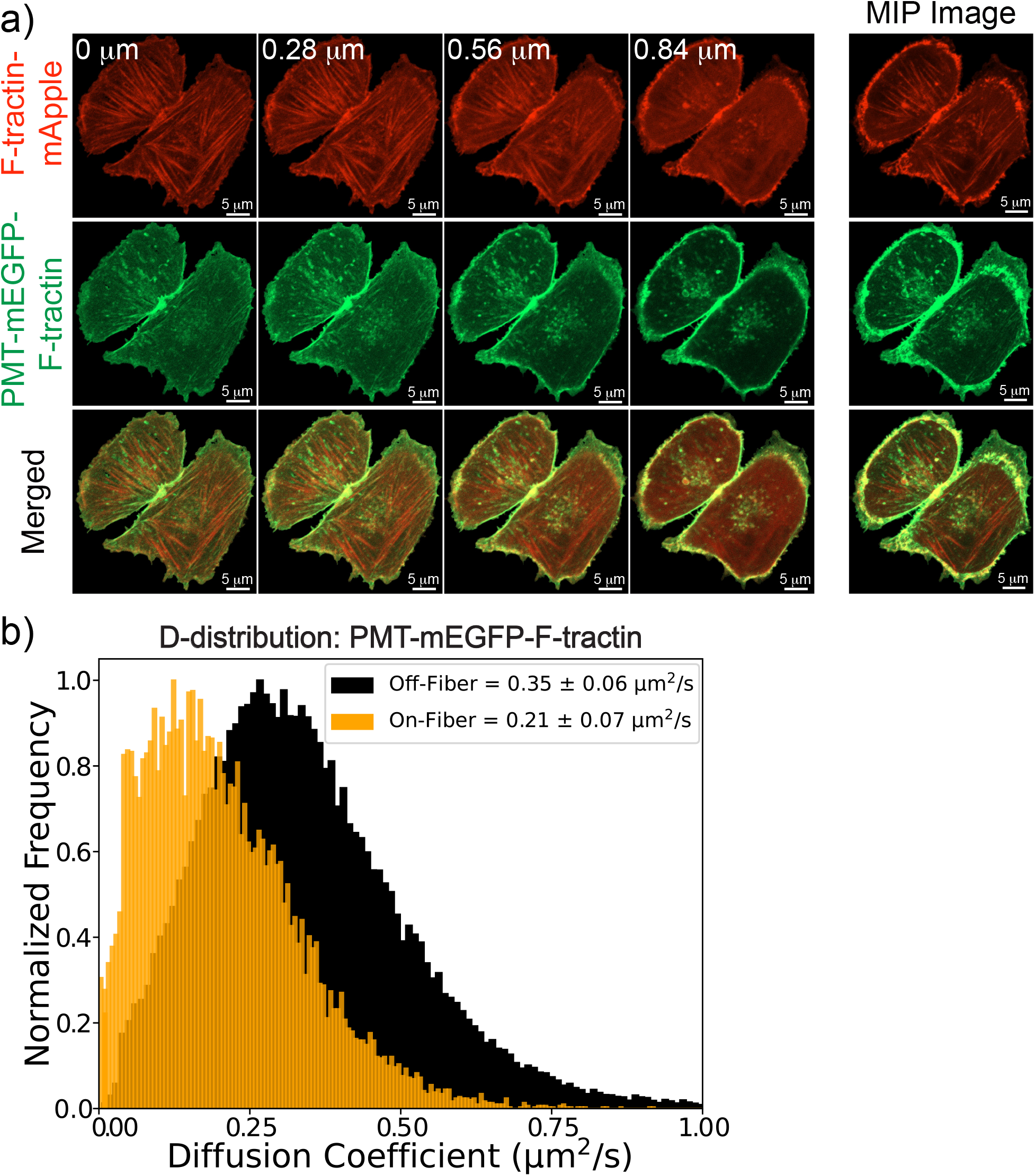
PMT-F-tractin as a CA-binding Structural and Dynamic Control. a) Z-stack confocal images at various depths and Maximum Intensity Projection (MIP) of F-tractin-mApple (top) and PMT-mEGFP-F-tractin(middle) illustrate distinct labeling patterns of the two probes in a representative CHO-K1 cell. The merged image (bottom) demonstrates preferential labeling of cortical actin (CA) by PMT-mEGFP-F-tractin, whereas F-tractin-mApple labels stress fibers in the deeper regions of the cell. b) Normalized frequency distribution of diffusion coefficients (D-distribution) across multiple cells expressing PMT-mEGFP-F-tractin reveals lower diffusion coefficients in regions containing fibers (on-fiber) compared to regions without fibers (off-fiber).

### EGFR dynamics and localization is independent of CA

To assess whether CA influences the organization and dynamics of EGFR, we analyzed spatial correlations in the intensity from the TIRF-averaged images and D from their D maps. We identified cell sections with fibers from the TIRF-averaged image and cropped corresponding areas from the D map for spatial autocorrelation analysis (78). This analysis measures how pixel values are correlated over varying distances, revealing pattern repetition in the input image. The presence of fibers in TIRF-averaged images of membrane proteins, regardless of their interaction with actin, facilitated the identification of fiber regions for EGFR and controls.

Given that PMT-mEGFP-F-tractin is a CA-binding probe, its D is expected to be influenced by the CA. Upon extracting the boxed region, as illustrated in the TIRF-averaged image (Figure 3a), we observed intensity correlation along the fibers. We also observed correlation along the fiber direction for D, confirming PMT-mEGFP-F-tractin’s D dependence on the CA.

**Figure 3:**
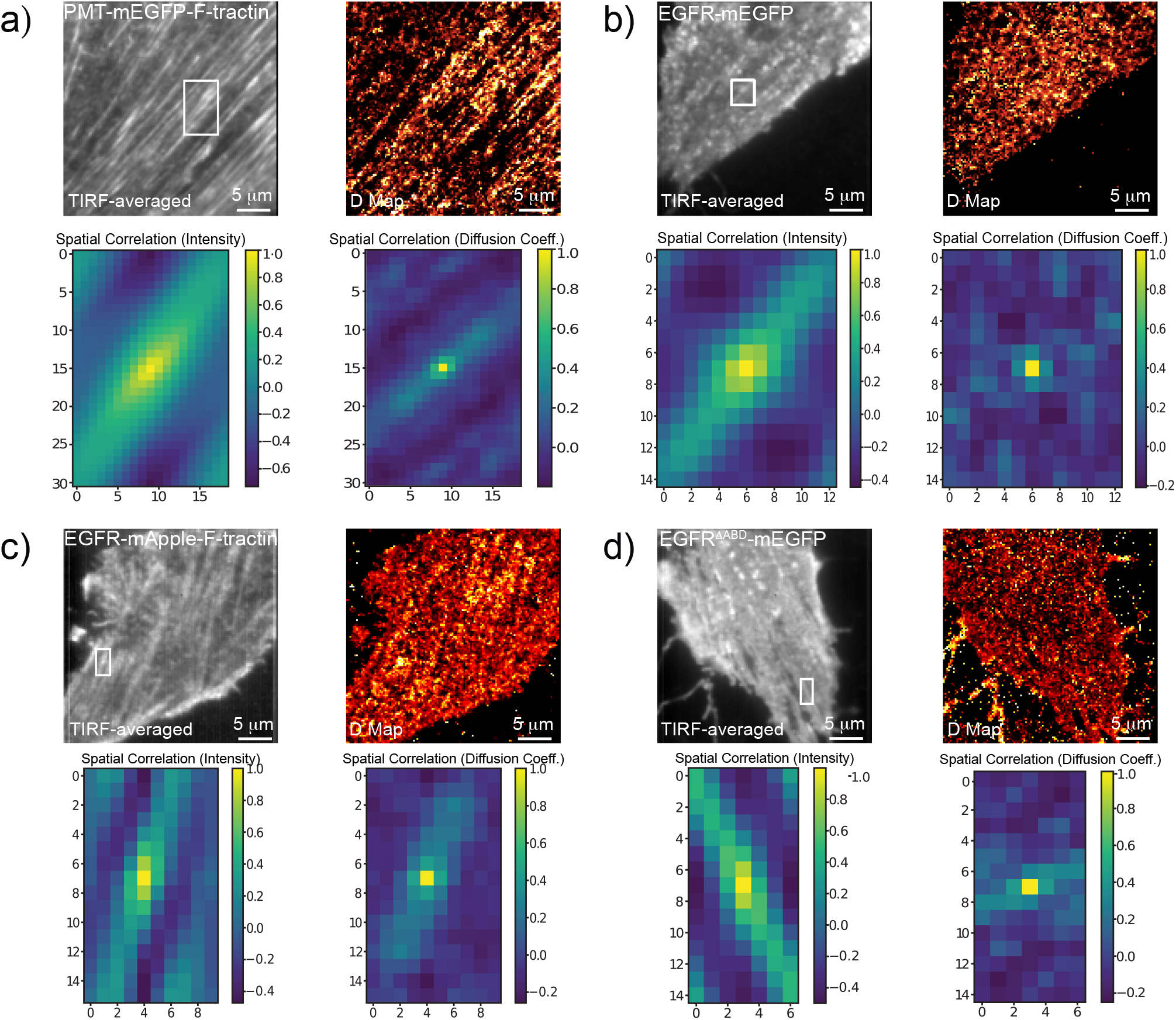
Spatial Correlation of Intensity and Diffusion Coefficient (D). Cell regions displaying fibers were identified in the TIRF-averaged image (boxed), and the region of interest (ROI) was cropped from the TIRF-averaged image and D-map. These ROIs were then used to generate spatial autocorrelation 2D maps with the same dimensions as the input files. The axes of the spatial autocorrelation maps indicate pixel numbers. Shown are the TIRF-averaged images, D maps, and their respective spatial autocorrelation maps for the boxed ROIs of (a) PMT-mEGFP-F-tractin, (b) EGFR-mEGFP, (c) EGFR-mApple-F-tractin, and (d) EGFR^ΔABD^-mEGFP.

EGFR-mEGFP, like other membrane proteins, exhibited fiber-like structures in its TIRF-averaged image, albeit at lower contrast (Figure 3b). Our dual-wavelength measurements of cells expressing EGFR-mEGFP and PMT-mApple-F-tractin showed that the fibers seen in the EGFR-mEGFP channel coincided with those labeled by PMT-mApple-F-tractin (Figure S2c). Spatial intensity correlation along the fiber regions from the two images confirms that the fibers observed with EGFR-mEGFP are actual actin fibers (Figure S2d). To investigate whether the D depends on these underlying fibers, we isolated the fiber-containing regions from EGFR-mEGFP TIRF-averaged image and the D map. While we observed spatial correlation in intensity along the fiber direction, there was no discernible correlation for D in the case of EGFR-mEGFP (Figure 3b). This suggests that CA does not influence the dynamics for EGFR-mEGFP.

Cells expressing EGFR-mApple-F-tractin display clusters specific to EGFR and fibers due to F-tractin binding. We selected a fiber region from an EGFR-mApple-F-tractin-expressing cell (Figure 3c). While the intensity exhibited a strong correlation along the fiber direction, we observed a subtle correlation pattern for D. EGFR-mApple-F-tractin shows some dependence on CA compared to EGFR-mEGFP but to a lesser extent than PMT-mEGFP-F-tractin. This is likely due to structural differences between the two F-tractin constructs, with PMT-mEGFP-F-tractin being closer to the membrane, while in EGFR-mApple-F-tractin, F-tractin is anchored at the end of the extended, unstructured C-terminal region of EGFR-mApple.

EGFR^ΔABD^-mEGFP, like EGFR-mEGFP, also shows the presence of fiber-like structures in its TIRF-averaged image (Figure 3d). When analyzed for spatial intensity correlation, we see correlations along the fiber direction. However, there is no observable correlation for D for the same fiber region, much like EGFR-mEGFP. Our spatial correlation results show that the localization and diffusion of EGFR-mEGFP and EGFR^ΔABD^-mEGFP remain independent of CA presence, unlike PMT-mEGFP-F-tractin and EGFR-mApple-F-tractin.

### EGFR shows no dynamics distinction in on-/off-fiber regions in resting or stress fiber inhibited state

To examine the influence of CA on EGFR diffusion, we analyzed the on- and off-fiber D-distributions in the cells’ resting state or following treatment with stress-fiber inhibitors. *D*_off_ and *D*_on_ signifies the average D values obtained from the *D*_off_ and *D*_on_ distributions (pooled from multiple cells) respectively. The *D*_off_ for PMT-mEGFP-F-tractin was higher than the *D*_on_ for both resting (Figure 4a) and stress fiber inhibited states (Figure 4b). In its resting state, PMT-mEGFP-F-tractin shows a *D*_off_ of 0.35 ± 0.06 µm^2^ s^−1^ and a lower *D*_on_ of 0.21 ± 0.07 µm^2^ s^−1^. In the stress fiber inhibited state, we see *D*_off_ of 0.42 ± 0.07 µm^2^ s^−1^ and *D*_on_ of 0.27 ± 0.08 µm^2^ s^−1^.

**Figure 4:**
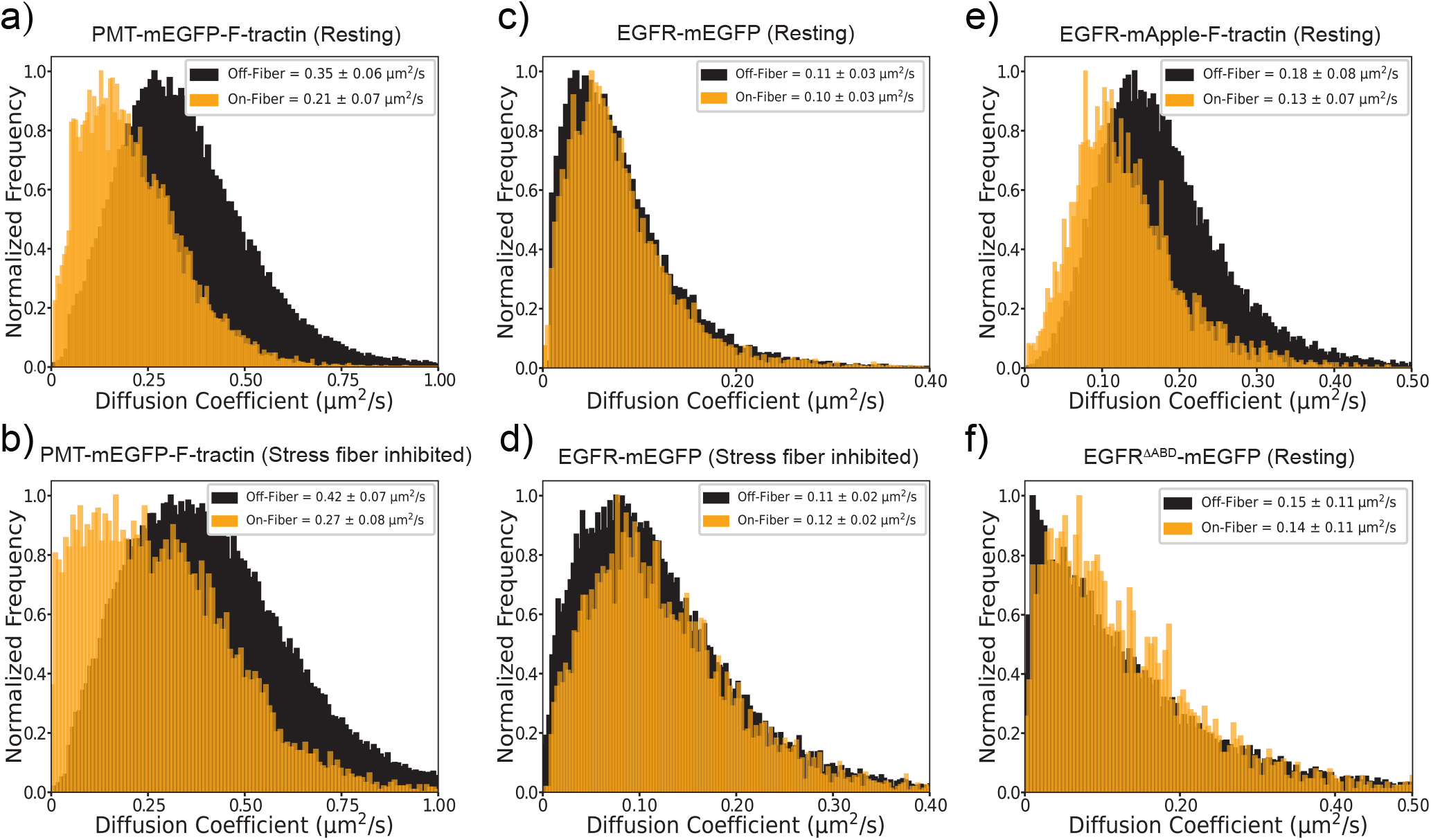
Normalized Frequency Distribution of Diffusion Coefficient (D) in on-/off-fiber regions for cells in either resting or stress-fiber inhibited state. On-fiber distributions are shown in orange and off-fiber distributions in black. Distributions are pooled from multiple cells expressing a) PMT-mEGFP-F-tractin (resting state), b) PMT-mEGFP-F-tractin (stress-fiber inhibited state), c) EGFR-mEGFP (resting state), d) EGFR-mEGFP (stress-fiber inhibited state), e) EGFR-mApple-F-tractin (resting state), and f) EGFR^ΔABD^-mEGFP (resting state). The inset shows the mean and SD of on-/off-fiber D.

After confirming the difference between *D*_on_ and *D*_off_ for PMT-mEGFP-F-tractin, we examined EGFR-mEGFP expressing cells with visible fibers. Pooled data from resting (Figure 4c) and stress fiber inhibited (Figure 4d) cells showed no discernible difference in *D*_on_ and *D*_off_ in either state. The average *D*_on_ or *D*_off_ value for EGFR-mEGFP remained consistent across both areas and states. In the resting state, *D*_off_ is 0.11 ± 0.03 µm^2^ s^−1^ and *D*_on_ is 0.10 ± 0.03 µm^2^ s^−1^. In the stress fiber inhibited state, *D*_off_ is 0.11 ± 0.02 µm^2^ s^−1^ and *D*_on_ is 0.12 ± 0.02 µm^2^ s^−1^, indicating that EGFR-mEGFP diffusion remains unaffected by CA or stress fibers, consistent with our prior findings (57).

For EGFR-mApple-F-tractin (Figure 4e) and EGFR^ΔABD^-mEGFP(Figure 4f), we selected cells showing fibers/fiber-like structures for each in their resting state. We observed a subtle increase in *D*_off_ (0.18 ± 0.08 µm^2^ s^−1^) for EGFR-mApple-F-tractin compared to *D*_on_ (0.13 ± 0.07 µm^2^ s^−1^). In contrast, EGFR^ΔABD^-mEGFP exhibited no difference in its *D*_off_ and *D*_on_, aligning with the trend observed for EGFR-mEGFP. The *D*_off_ is 0.15 ± 0.11 µm^2^ s^−1^ and *D*_on_ is 0.14 ± 0.11 µm^2^ s^−1^. This suggests that the dynamics of EGFR-mEGFP and EGFR^ΔABD^-mEGFP in the resting state are independent of CA, but EGFR can be made CA-sensitive by adding F-tractin to its C-terminus. The average values for *D*_off_ and *D*_on_ for the different constructs pooled from multiple cells can be found in the Supplementary Table S1.

We performed simultaneous dual-wavelength measurements to label the CA fibers with PMT-mEGFP-F-tractin and measure the dynamics of EGFR-mApple or EGFR-mApple-F-tractin. We observed that the fibers labeled by PMT-mEGFP-F-tractin are also seen in the EGFR-mApple-F-tractin channel (Figure S3a). When the marked cell region spanning across a fiber is extracted from the TIRF-averaged images of PMT-mEGFP-F-tractin, EGFR-mApple-F-tractin, and the corresponding D-map of EGFR-mApple-F-tractin, we observe a strong correlation in the intensity of PMT-mEGFP-F-tractin and EGFR-mApple-F-tractin, with a Pearson’s correlation coefficient, R, of 0.94 (Figure S3b). The D of EGFR-mApple-F-tractin and the intensity of PMT-mEGFP-F-tractin across the marked region show moderate anti-correlation with an R-value of -0.69 (Figure S3c). This confirms that EGFR-mApple-F-tractin interacts with CA, and its dynamics are influenced by this interaction. In the dual-channel experiment with EGFR-mApple and PMT-mEGFP-F-tractin, we observed fibers in the EGFR-mApple channel as we have observed with single-channel measurements of EGFR-mEGFP (Figure S3d). Extracting a region spanning the fiber seen in the PMT-mEGFP-F-tractin channel and correlating the intensity of PMT-mEGFP-F-tractin with EGFR-mApple shows a positive correlation with an R-value of 0.84 (Figure S3e). The intensity peaks of PMT-mEGFP-F-tractin and EGFR-mApple coincide, confirming that the fibers seen in the EGFR-mApple channel are the same fibers labeled by the CA probe, PMT-mEGFP-F-tractin. Correlating the D of EGFR-mApple with the intensity of PMT-mEGFP-F-tractin shows no observable correlation, with an R-value of 0.10 (Figure S3f). These dual-wavelength measurements show that EGFR dynamics are not influenced by CA.

### Actin-organization perturbations show limited influence on EGFR and Δ ABD-EGFR dynamics

The cells expressing EGFR and controls were treated with Latrunculin-A (Lat-A) and Jasplakinolide (Jas), which affect the cytoskeleton by disrupting F-actin and stabilizing/promoting actin polymerization, respectively. The dynamics of PMT-mEGFP-F-tractin show significant variations following drug treatments (Figure 5a). In the case of Jas treatment, D decreases from 0.29 ± 0.10 µm^2^ s^−1^ to 0.22 ± 0.10 µm^2^ s^−1^. Conversely, Lat-A treatment results in an increase in the D from 0.29 ± 0.10 µm^2^ s^−1^ to 0.43 ± 0.10 µm^2^ s^−1^. For EGFR-mEGFP, actin perturbation does not elicit substantial changes in its dynamics, with D around 0.14 ± 0.04 µm^2^ s^−1^ under both conditions (Figure 5b). The D of EGFR-mApple-F-tractin does not show significant change after Jas treatment (0.22 ± 0.06 µm^2^ s^−1^ from 0.23 ± 0.10 µm^2^ s^−1^), while a slight increase is observed following Lat-A treatment (0.30 ± 0.09 µm^2^ s^−1^ from 0.23 ± 0.10 µm^2^ s^−1^) (Figure 5c). The significant D difference obtained post-Lat-A treatment arises from the binding of F-tractin to the actin framework. However, the D-differences obtained for EGFR-mApple-F-tractin are not as pronounced as PMT-mEGFP-F-tractin. This observation might be due to the structural difference between the two F-tractin constructs. PMT-mEGFP-F-tractin binds to membrane-proximal CA due to membrane tethering by PMT, while EGFR-mApple-F-tractin interacts with the CA via F-tractin anchored at the end of the extended, unstructured C-terminus of EGFR. The dynamics of EGFR^ΔABD^-mEGFP remain largely unaffected, exhibiting a diffusion coefficient of around 0.16 ± 0.04 µm^2^ s^−1^ under each condition (Figure 5d), which is consistent with the trend observed for EGFR-mEGFP. These results suggest that while the actin cytoskeleton significantly influences the dynamics of PMT-mEGFP-F-tractin, it has a limited impact on EGFR-mEGFP and EGFR^ΔABD^-mEGFP. EGFR dynamics are largely independent of actin perturbations unless EGFR is modified to directly interact with the actin network. The D obtained for different constructs under these actin-perturbing cell treatments pooled from multiple cells are tabulated in Table S2.

**Figure 5:**
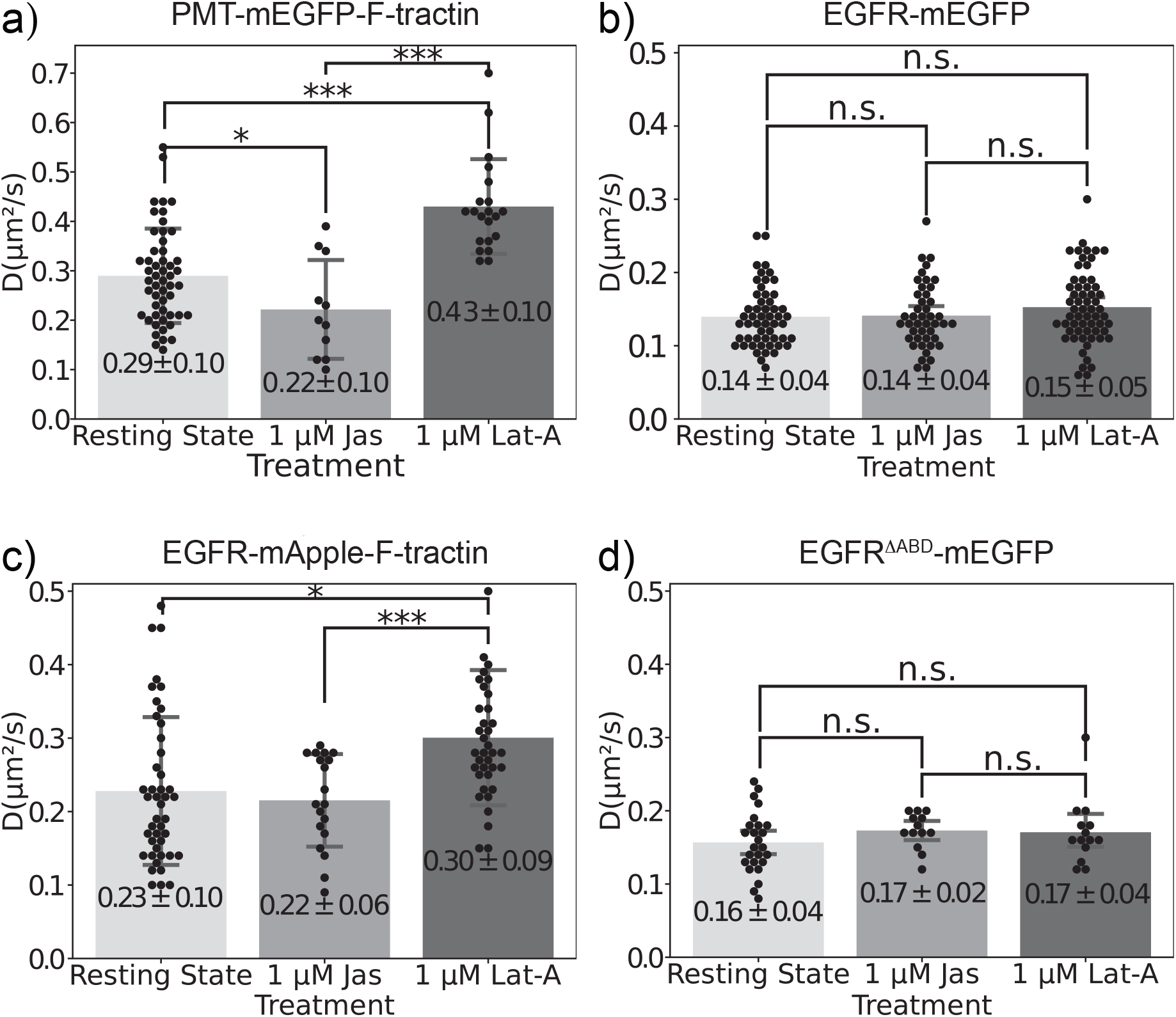
Effect of actin-perturbing drugs on the dynamics of. a) PMT-mEGFP-F-tractin, b) EGFR-mEGFP, c) EGFR-mApple-F-tractin, and d) EGFR^ΔABD^-mEGFP. Data points represent individual measurements, with bars showing mean ± SD of D. Significance was calculated using ANOVA with post-hoc tests: *** p < 0.001, * p < 0.05, and n.s.: non-significant.

It is noteworthy that EGFR-mApple-F-tractin exhibits a higher D than EGFR-mEGFP, despite the former having an additional F-tractin anchor. This discrepancy likely stems from differences in the fluorophores used for labeling these proteins. Red fluorescent proteins like mApple exhibit lower fluorescence intensity and increased photobleaching compared to green fluorescent proteins such as mEGFP. In FCS, this can lead to so-called cryptic photobleaching, i.e., the fluorescent tag bleaches before the particle transits the observation volume, leading to an overestimation of D (79). This is consistent with the fact that mApple-tagged constructs show higher D values than their mEGFP-tagged counterparts, as can be seen with the membrane proteins, PMT (Figure S4a) and PMT-F-tractin (Figure S4b). Individual D values for cells expressing PMT-mEGFP/PMT-mApple/PMT-mEGFP-F-tractin/PMT-mApple-F-tractin(Table S3) show that PMT-mApple and PMT-mApple-F-tractin exhibit a higher D than PMT-mEGFP and PMT-mEGFP-F-tractin, respectively. Thus, our analysis focuses on relative differences in D for the same construct across different treatment conditions or cell states rather than comparing D values across different proteins.

### EGF-stimulated EGFR does not directly interact with CA

To understand the EGFR/CA interaction in the receptor’s activated state, we stimulated EGFR and controls with its extracellular ligand, EGF. We pooled the D-data from multiple cells in their resting state, followed by stimulation with EGF for receptor activation, and then Lat-A treatment to depolymerize the actin framework. This approach allowed us to measure changes in D after stimulation and subsequent actin disruption. These changes served as the test to assess activated EGFR’s dependency on CA and to investigate if the putative ABD becomes functional post-activation and whether EGFR then interacts or directly binds to CA.

In the case of EGFR-mEGFP, post-EGF stimulation, the average D decreased by 60%, from a resting state D of 0.14 ± 0.04 µm^2^ s^−1^ to an activated state D of 0.06 ± 0.03 µm^2^ s^−1^ (Figure 6a). EGF binding promotes EGFR oligomerization, which is essential for the activation of the receptor’s intrinsic kinase activity, resulting in the observed decrease in D post-stimulation. When cells were treated with Lat-A post-stimulation, there was almost no change in the average D value, which remained at 0.07 ± 0.03 µm^2^ s^−1^. EGFR-mEGFP in its resting state also does not show any significant difference post Lat-A treatment (Figure 5b), which is also observed with activated EGFR-mEGFP.

**Figure 6:**
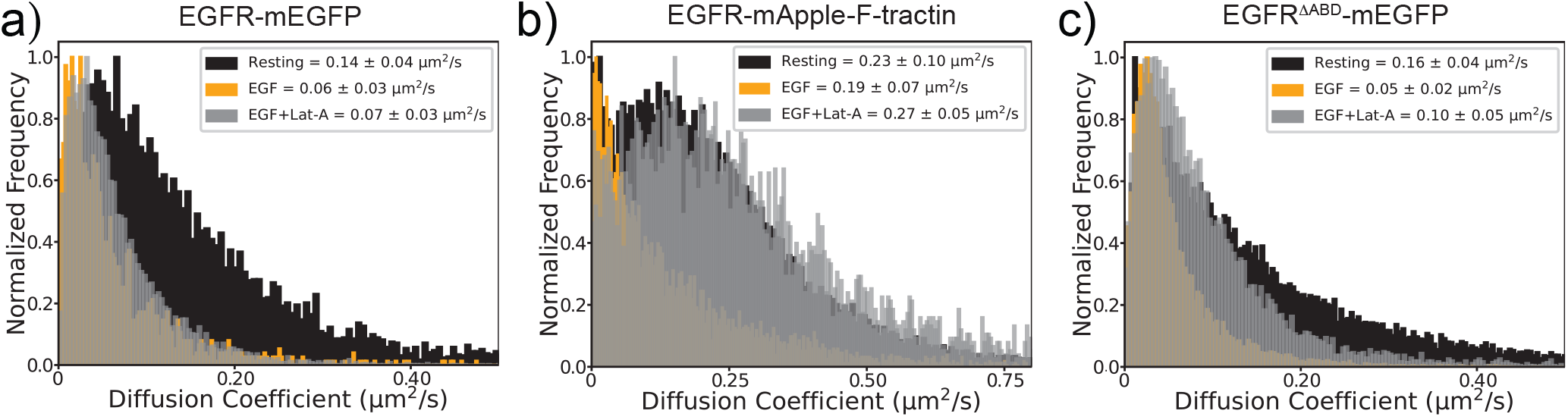
Normalized frequency distribution of Diffusion Coefficient (D) post actin disruption in the EGF-induced activated state for a) EGFR-mEGFP b) EGFR-mApple-F-tractin and c) EGFR^ΔABD^-mEGFP. The inset shows mean ± SD of the D. Distribution colors: black for resting state, orange for post-EGF-stimulation, and grey for post-EGF-stimulation, followed by Lat-A treatment.

For EGFR-mApple-F-tractin, the resting state showed an average D of 0.23 ± 0.10 µm^2^ s^−1^. D decreased post-EGF stimulation to 0.19 ± 0.07 µm^2^ s^−1^ (Figure 6b). Lat-A treatment in both resting and activated states for EGFR-mApple-F-tractin showed a significant increase in D. In the resting state, D increased by 30%, from 0.23 ± 0.10 µm^2^ s^−1^ to 0.30 ± 0.09 µm^2^ s^−1^(Figure 5b).

In its activated state, D again increased by almost 40%, from 0.19 ± 0.07 µm^2^ s^−1^ to 0.27 ± 0.05 µm^2^ s^−1^ post-Lat-A treatment (Figure 6b).

EGFR^ΔABD^-mEGFP exhibited a reduced D post-EGF stimulation, decreasing from 0.16 ± 0.04 µm^2^ s^−1^ to 0.05 ± 0.02 µm^2^ s^−1^ after stimulation (Figure 6c). Post-activation, Lat-A induced a slight increase in the average D to 0.10 ± 0.05 µm^2^ s^−1^, with a subtle rightward shift in distribution.

Together, Lat-A treatment-induced differences in the D for EGFR constructs before and after EGF stimulation are tabulated in Table S4. The decrease in D post-EGF stimulation was consistently observed for EGFR-mEGFP, EGFR^ΔABD^-mEGFP, and EGFR-mApple-F-tractin, attributable to EGFR’s ligand-induced oligomerization and activation. Actin disruption had minimal effect on the dynamics of activated EGFR-mEGFP, suggesting its independence from the CA. In contrast, resting and activated EGFR-mApple-F-tractin’s altered dynamics to Lat-A implies its direct CA dependence. Notably, EGFR-mEGFP did not demonstrate direct binding or interaction with CA, even in its activated state.

## GENERAL DISCUSSION

In this SRRF’n’TIRF study, we investigated the influence of cortical actin (CA) on EGFR in its resting and activated states. The combination of SRRF and ITIR-FCS allows us to simultaneously determine super-resolved structures and millisecond-scale molecular dynamics from the same dataset. EGFR-actin interaction studied previously using SRRF’n’TIRF has shown that the static cytoskeleton indirectly affects the dynamics and organization of EGFR by influencing the membrane (57). However, the actin signal obtained was a mix of membrane-proximal CA and stress fibers from the cytoplasm due to the lack of a CA-specific probe, making it unclear whether EGFR directly interacts with CA. To address this, we constructed PMT-F-tractin as a CA-targeting probe, inspired by (62).

TIRF-averaged images, being diffraction-limited, contain mixed signals from fiber and surrounding regions, causing ambiguity in the exact positioning of on-fiber pixels. In contrast, the SRRF reconstruction offers a super-resolved depiction of the fibers, enabling precise identification of on-fiber pixels and allowing us to create binary on-fiber masks. As SRRF creates additional artifacts, we corrected these by intensity thresholding to maximize fiber retention before generating the on-fiber mask (57). TIRF images, due to their diffraction-limited resolution, identify not only pixels directly on top of actin fibers but also neighboring pixels with signals from off-fiber regions. Thus, we used off-fiber pixels identified in TIRF to create a binary off-fiber mask that identifies pixels that are not influenced by any signal from fibers. These masks overlaid on the D-Map allowed us to isolate and analyze on-/off-fiber D maps and their distributions, avoiding intermediate regions of mixed diffusion coefficients, which helped us to clearly distinguish and analyze on- and off-fiber diffusion behavior.

PMT-mEGFP-F-tractin showed expected diffusion differences in regions with varying CA density, exhibiting slower diffusion in on-fiber regions and faster diffusion in off-fiber regions. Unlike F-tractin and Lifeact, which bind broadly to CA and stress fibers, PMT-F-tractin consistently targeted the CA network. This made PMT-mEGFP-F-tractin a good CA-binding structural and dynamic probe.

We consistently observed actin fibers in the TIRF-averaged images of membrane proteins, which was surprising for non-actin-binding membrane proteins like GFP-GPI, PMT-mEGFP, and EGFR^ΔABD^-mEGFP. This phenomenon suggests that actin cytoskeleton-induced membrane morphology may play a role, which would be consistent with recent reports that membrane proteins can accumulate in regions of increased membrane curvature (80–84). The visibility of these fibers decreases with increased penetration depth, supporting the idea that these structures are seen at the membrane (albeit at a lower contrast than seen with actin-binding proteins) and are, thus, a membrane property. Dual wavelength measurements on cells expressing PMT-mApple-F-tractin and EGFR-mEGFP confirmed that the fiber-like structures observed with EGFR-mEGFP are indeed actin fibers also labeled by PMT-mApple-F-tractin. This allowed us to compare EGFR and its controls’ dependence and interaction with the CA even without specifically labeling it with PMT-F-tractin and compromising on the SNR. Our spatial correlation analysis, on-/off-fiber D-distributions, and actin perturbing drug-treatment studies consistently show that resting-state EGFR-mEGFP dynamics and organization are independent of CA density. Even in the EGF-stimulated, activated state, EGFR-mEGFP shows no direct interaction with the CA.

Anchoring an actin-binding peptide to the C-terminus of EGFR renders it dependent on CA. EGFR-mEGFP and EGFR^ΔABD^-mEGFP show similar diffusion coefficient distributions and organization, suggesting that the putative ABD has limited functionality in both resting and activated states. EGFR-mApple-F-tractin shows more CA dependence than the other two EGFR constructs, aligning closely in behavior with the CA-binding probe, PMT-mEGFP-F-tractin. Overall, our findings indicate that EGFR-mEGFP does not directly bind, or at the very least not strongly bind, to CA in either the resting or activated state.

## AUTHOR CONTRIBUTIONS

TW conceived of the study. SP performed experiments, analysed the data, and wrote the first draft, all authors contributed to the final version of the manuscript.

## ACKNOWLEDGMENTS

TW gratefully acknowledges funding from the Singapore Ministry of Education (MOE2016-T3-1-005). SP is a recipient of a research scholarship from the National University of Singapore.

