## Supplementary Materials for "EGFR does not directly interact with cortical actin: A SRRF’n’TIRF Study"

**SUPPLEMENTARY MATERIAL**

70 nm	80 nm	90 nm	100 nm	120 nm	140 nm

a)

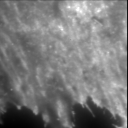

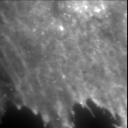

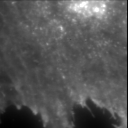

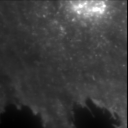

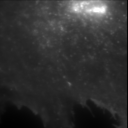

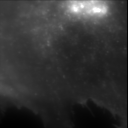

b)

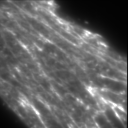

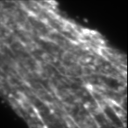

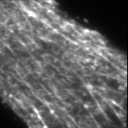

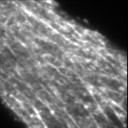

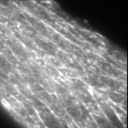

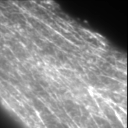

c)

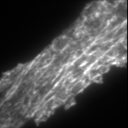

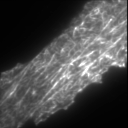

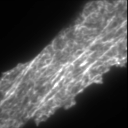

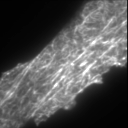

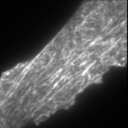

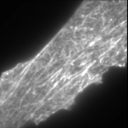

Figure S1: TIRF-averaged images of membrane proteins at different penetration depths show the loss in contrast of actin fibers as the penetration depth increases for non-actin binding proteins. Panels:(a) GFP-GPI, (b) F-tractin-mEGFP, (c) PMT-mEGFP- F-tractin. Scale bar: 5 µm.

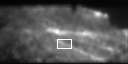

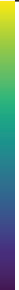

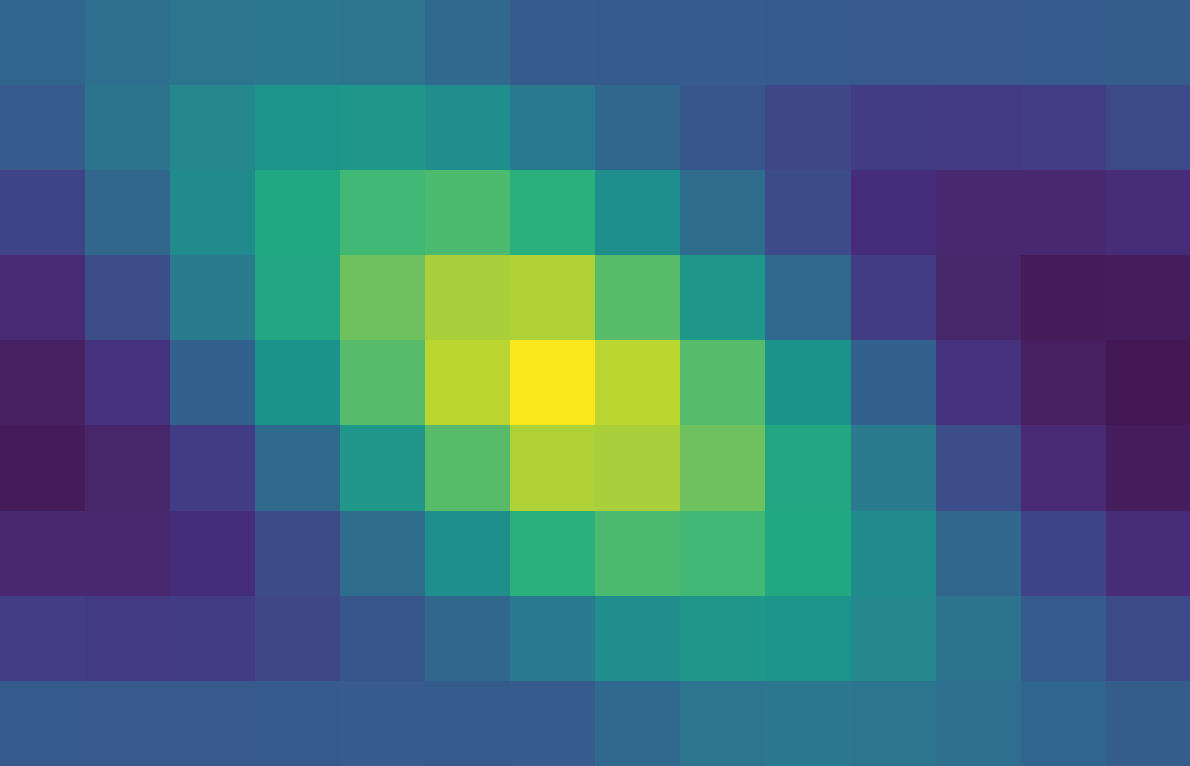

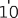

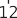

a) PMT-mApple-F-tractin

b)

Spatial Intensity Correlation (PMT-mApple-F-tractin)

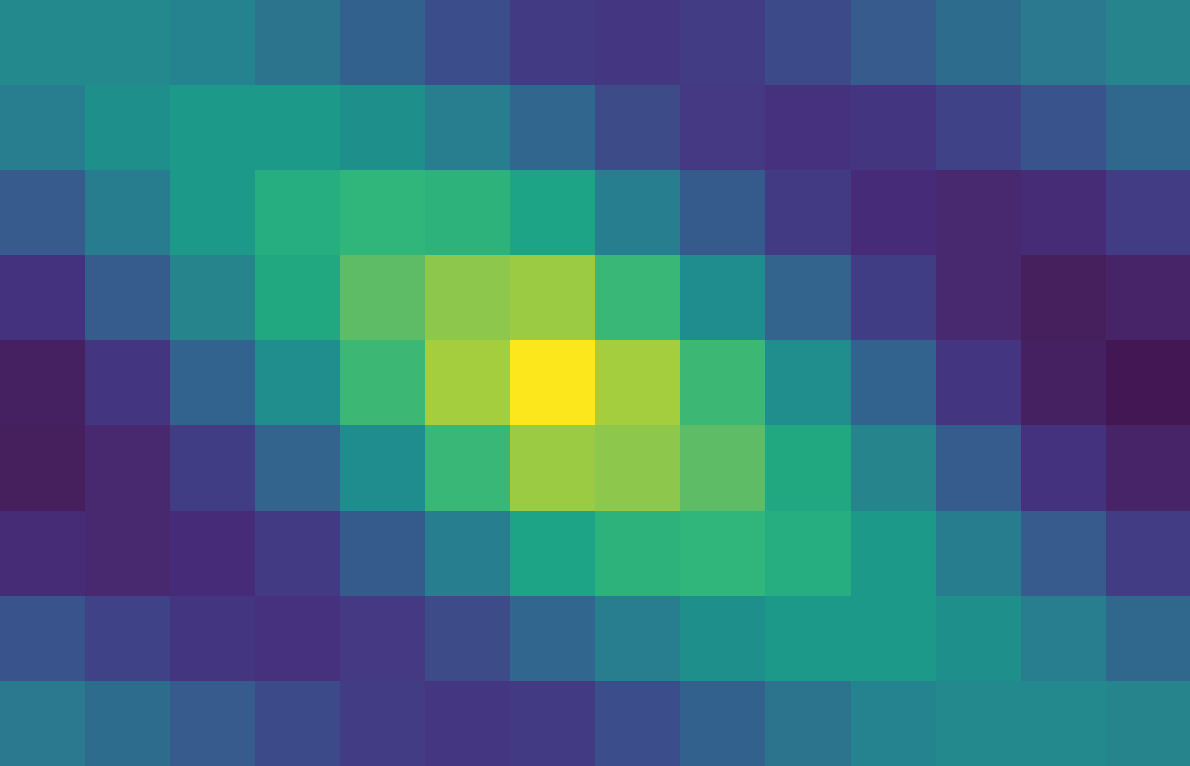

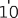

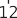

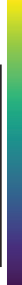

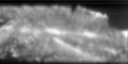

GFP-GPI

Spatial Intensity Correlation (GFP-GPI)

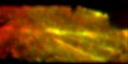

Merged

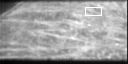

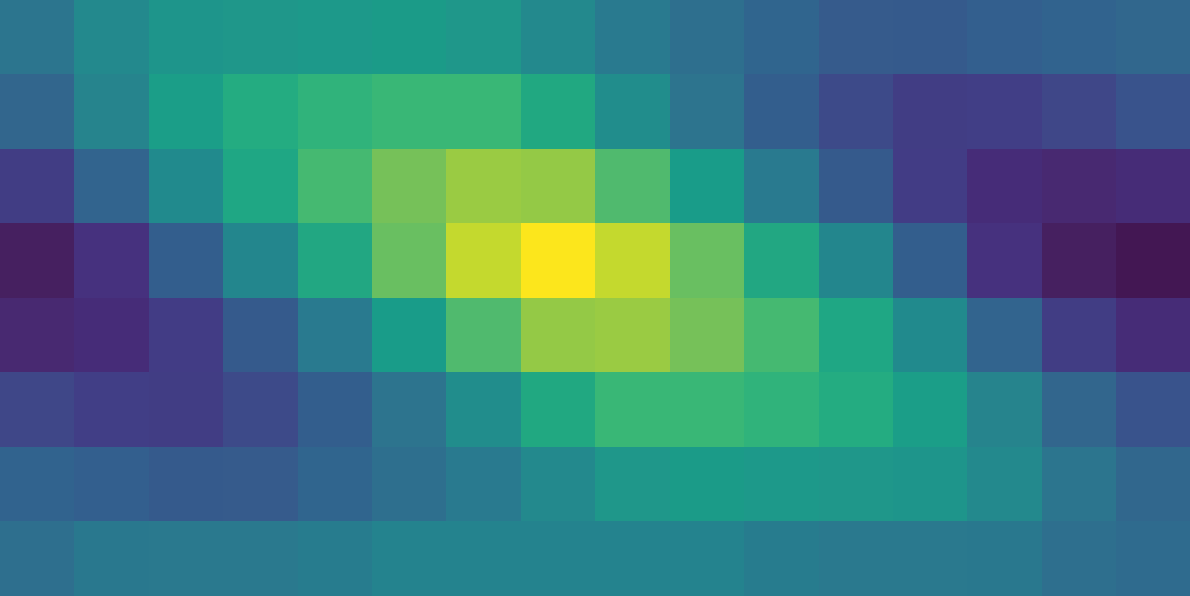

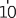

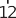

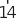

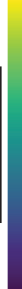

c) PMT-mApple-F-tractin

d)

Spatial Intensity Correlation (PMT-mEGFP-F-tractin)

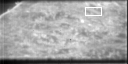

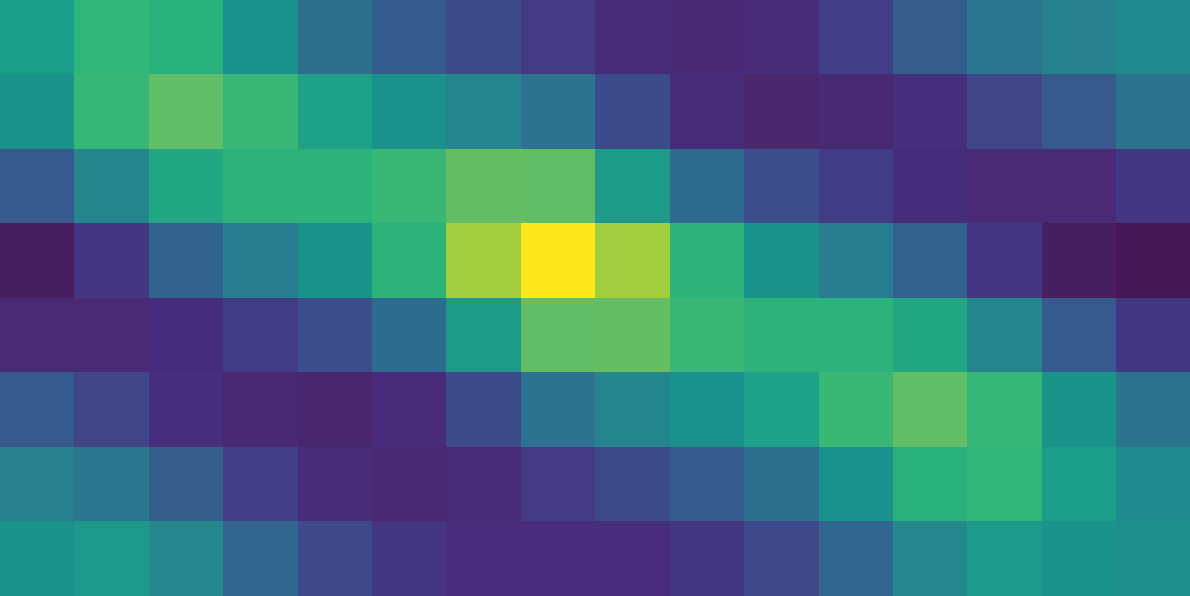

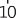

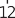

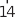

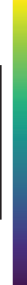

EGFR-mEGFP

Spatial Intensity Correlation (EGFR-mEGFP)

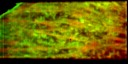

Merged

Figure S2: Simultaneous dual-wavelength imaging of cells expressing: a) PMT-mApple-F-tractin and GFP-GPI, or c) PMT- mApple-F-tractin and EGFR-mEGFP. Left: PMT-mApple-F-tractin; Middle: GFP-GPI or EGFR-mEGFP; Right: Merged overlay (red: PMT-mApple-F-tractin, green: GFP-GPI or EGFR-mEGFP). b,d) Spatial intensity correlations of boxed regions showing fiber alignment. Autocorrelation map axes in pixels. Scale bar: 5 µm.

Table S1: Average *𝐷*_off_ and *𝐷*_on_ values for different proteins pooled from multiple cells in resting or stress fiber inhibited state

| **Protein** | **Cell State** | *𝐷***_off_ (**µm2 s−1**)** | *𝐷***_on_ (**µm2 s−1**)** | **Number of Cells** |
| --- | --- | --- | --- | --- |
| PMT-mEGFP-F-tractin | Resting | 0.35 ± 0.06 | 0.21 ± 0.07 | 11 |
| PMT-mEGFP-F-tractin | Stress fiber inhibited | 0.42 ± 0.07 | 0.27 ± 0.08 | 13 |
| EGFR-mEGFP | Resting | 0.11 ± 0.03 | 0.10 ± 0.03 | 9 |
| EGFR-mEGFP | Stress fiber inhibited | 0.11 ± 0.02 | 0.12 ± 0.02 | 13 |
| EGFR-mApple-F-tractin | Resting | 0.18 ± 0.08 | 0.13 ± 0.07 | 4 |
| EGFRΔABD-mEGFP | Resting | 0.15 ± 0.11 | 0.14 ± 0.11 | 4 |

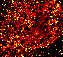
a)

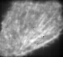

EGFR-mApple-F-tractin

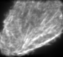

PMT-mEGFP-F-tractin

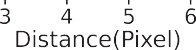

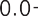

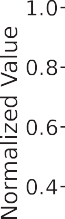

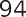

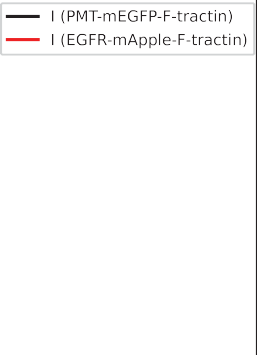

b)

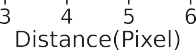

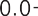

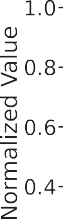

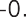

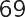

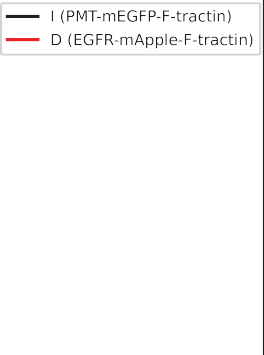

c)

d)

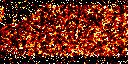
f)

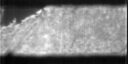

EGFR-mApple

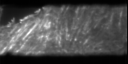

PMT-mEGFP-F-tractin

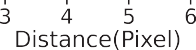

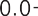

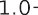

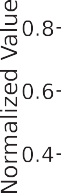

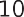

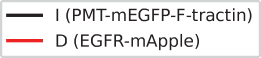

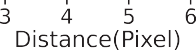

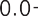

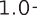

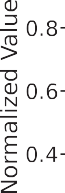

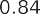

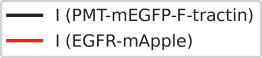

e)

Figure S3: Simultaneous dual-wavelength imaging of cells expressing (a) PMT-mEGFP-F-tractin and EGFR-mApple-F-tractin or (d) PMT-F-tractin-mEGFP and EGFR-mApple. (a,d) Left: PMT-F-tractin-mEGFP; Middle: EGFR construct; Right: D-map of EGFR construct. Black lines indicate regions for correlation analysis. (b,e) Intensity correlation along marked lines (PMT-F-tractin: black, EGFR construct: red). (c,f) Correlation between PMT-F-tractin intensity (black) and EGFR construct diffusion coefficient (red). R values indicated. Scale bar: 5 µm.

Table S2: Diffusion coefficients (D) pooled from multiple cells after treatment with actin-perturbing drugs for different protein constructs

| **Cell Treatment**  **Protein** | **No Treatment (Resting)** | | **Jasplakinolide (Jas)** | | **Latrunculin-A (Lat-A)** | |
| --- | --- | --- | --- | --- | --- | --- |
|  | **D (**µm^2^ s^−1^**)** | **No. of Cells** | **D (**µm^2^ s^−1^**)** | **No. of Cells** | **D (**µm^2^ s^−1^**)** | **No. of Cells** |
| PMT-mEGFP-F-tractin | 0.29 ± 0.10 | 52 | 0.22 ± 0.10 | 11 | 0.43 ± 0.10 | 21 |
| EGFR-mEGFP | 0.14 ± 0.04 | 53 | 0.14 ± 0.04 | 44 | 0.15 ± 0.05 | 60 |
| EGFR-mApple-F-tractin | 0.23 ± 0.10 | 44 | 0.22 ± 0.06 | 19 | 0.30 ± 0.09 | 37 |
| EGFRΔABD-mEGFP | 0.16 ± 0.04 | 25 | 0.17 ± 0.02 | 13 | 0.17 ± 0.04 | 14 |

a)	D Values for PMT-mEGFP/-mApple	b)	D Values for PMT-mEGFP/-mApple-F-tractin

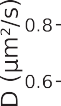

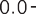

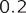

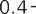

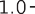

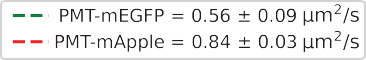

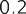

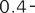

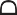

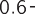

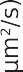

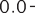

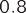

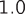

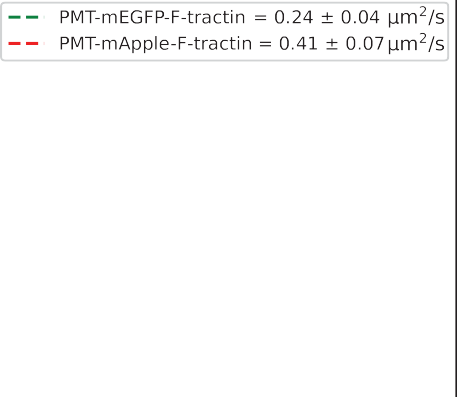

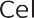

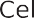

Figure S4: Diffusion coefficient (D) for cells expressing (a) PMT-mEGFP (green) or PMT-mApple (red), (b) PMT-mEGFP-F- tractin (green) or PMT-mApple-F-tractin (red) across six different cells. The inset shows the mean and standard deviation of the D values for mEGFP/mApple-tagged proteins across the six cells.

Table S3: Diffusion Coefficient (D) Values for mEGFP and mApple-tagged proteins across six different cells

| **Cell No.** | **PMT-mEGFP** | **PMT-mApple** | **PMT-mEGFP-F-tractin** | **PMT-mApple-F-tractin** |
| --- | --- | --- | --- | --- |
|  | **Diffusion Coefficient (µm^2^/s)** | | | |
| 1 | 0.59 ± 0.30 | 0.85 ± 0.28 | 0.29 ± 0.14 | 0.36 ± 0.22 |
| 2 | 0.62 ± 0.21 | 0.88 ± 0.28 | 0.18 ± 0.09 | 0.35 ± 0.19 |
| 3 | 0.40 ± 0.30 | 0.78 ± 0.35 | 0.20 ± 0.10 | 0.54 ± 0.22 |
| 4 | 0.68 ± 0.36 | 0.87 ± 0.29 | 0.27 ± 0.11 | 0.37 ± 0.21 |
| 5 | 0.53 ± 0.31 | 0.83 ± 0.34 | 0.27 ± 0.14 | 0.38 ± 0.23 |
| 6 | 0.51 ± 0.30 | 0.84 ± 0.30 | 0.26 ± 0.10 | 0.48 ± 0.23 |

Table S4: Diffusion coefficients (D) in untreated and Lat-A treated cells for EGFR constructs with or without EGF stimulation pooled from multiple cells

| **Protein** | **-EGF** | | | | **+EGF** | | | |
| --- | --- | --- | --- | --- | --- | --- | --- | --- |
|  | **Untreated** | | **Lat-A treated** | | **Untreated** | | **Lat-A treated** | |
|  | **D (**µm^2^ s^−1^**)** | **No. of Cells** | **D (**µm^2^ s^−1^**)** | **No. of Cells** | **D (**µm^2^ s^−1^**)** | **No. of Cells** | **D (**µm^2^ s^−1^**)** | **No. of Cells** |
| EGFR-mEGFP | 0.14 ± 0.04 | 53 | 0.15 ± 0.05 | 60 | 0.06 ± 0.03 | 6 | 0.07 ± 0.03 | 10 |
| EGFR-mApple-F-tractin | 0.23 ± 0.10 | 44 | 0.30 ± 0.09 | 37 | 0.19 ± 0.07 | 6 | 0.27 ± 0.05 | 10 |
| EGFRΔABD-mEGFP | 0.16 ± 0.04 | 25 | 0.17 ± 0.04 | 14 | 0.05 ± 0.02 | 13 | 0.10 ± 0.05 | 16 |
